# Identification, characterization, and application of a highly sensitive lactam biosensor from *Pseudomonas putida*

**DOI:** 10.1101/700484

**Authors:** Mitchell G. Thompson, Allison N. Pearson, Jesus F. Barajas, Pablo Cruz-Morales, Nima Sedaghatian, Zak Costello, Megan E. Garber, Matthew R. Incha, Luis E. Valencia, Edward E. K. Baidoo, Hector Garcia Martin, Aindrila Mukhopadhyay, Jay D. Keasling

## Abstract

Caprolactam is an important polymer precursor to nylon traditionally derived from petroleum and produced on a scale of 5 million tons per year. Current biological pathways for the production of caprolactam are inefficient with titers not exceeding 2 mg/L, necessitating novel pathways for its production. As development of novel metabolic routes often require thousands of designs and result in low product titers, a highly sensitive biosensor for the final product has the potential to rapidly speed up development times. Here we report a highly sensitive biosensor for valerolactam and caprolactam from *Pseudomonas putida* KT2440 which is >1000x more sensitive to exogenous ligand than previously reported sensors. Manipulating the expression of the sensor *oplR* (PP_3516) substantially altered the sensing parameters, with various vectors showing K_d_ values ranging from 700 nM (79.1 μg/L) to 1.2 mM (135.6 mg/L). Our most sensitive construct was able to detect *in vivo* production of caprolactam above background at ~6 μg/L. The high sensitivity and range of OplR is a powerful tool towards the development of novel routes to the biological synthesis of caprolactam.

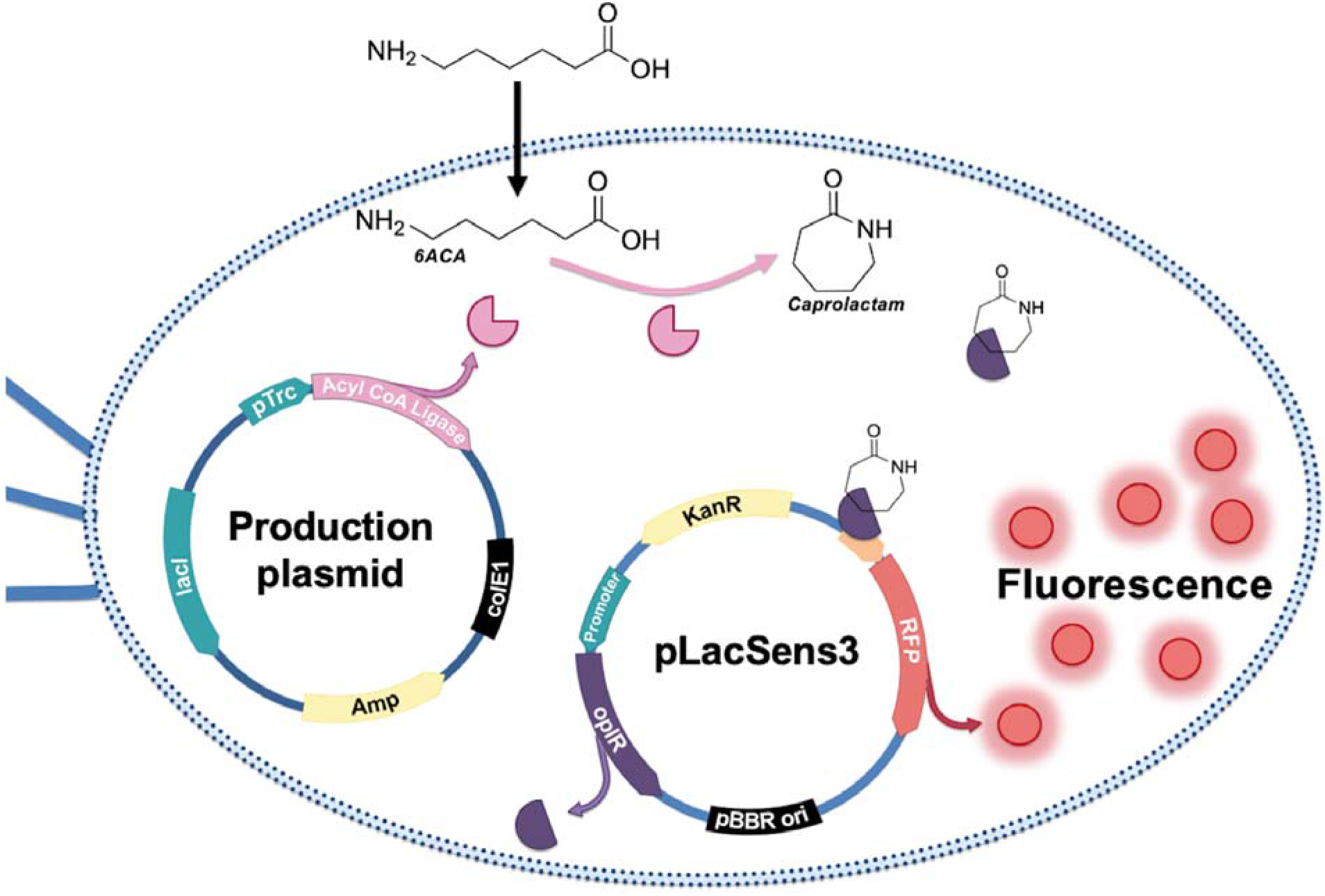

## INTRODUCTION

Caprolactam is an important chemical precursor to the polymer nylon 6, with a global demand approximately 5 million tons per year^1^. Currently the majority of caprolactam is synthesized from cyclohexanone, which in turn derived from petroleum^2^. In addition to being inherently unsustainable, the chemical process to synthesize caprolactam requires toxic reagents and produces unwanted byproducts such as ammonium sulfate^2^. Multiple attempts have been made to produce caprolactam biologically, however the highest titers achieved to date are no greater than 1-2 mg/L^1,3^. All current published strategies to produce caprolactam *in vivo* rely on the cyclization of 6-aminocaproic acid (6ACA) via a promiscuous acyl-coA ligase^1,3^. While similar strategies to make C4 butyrolactam and C5 valerolactam produce gram per liter titers of each, it is thought that both the entropy and enthalpy properties of 7-membered ring formation of caprolactam present an inherent barrier to the cyclization of 6ACA^4,5^. Renewable caprolactam production has been further hampered by poor yields of 6ACA *in vivo^6^*, as well as challenges in controlling the chain length of ω-amino acids. Clearly, novel routes to a renewable biological production of caprolactam are needed^7^.

Genetically encoded biosensors can accelerate metabolic engineering efforts in many ways, the foremost of which is the ability to rapidly screen for desirable phenotypes beyond the throughput of analytical chemistry^8^. Multiple papers have reported transcription factors or riboswitches that respond to lactams with varying degrees of sensitivity and specificity^4,9,10^. One feature that unifies currently available lactam biosensors is that lactams are not the native ligand for any of the corresponding biosensor systems. Zhang et al. used ChnR from *Acinetobacter* sp. Strain NCIMB 9871 to sense multiple lactams with all ligands tested having a K_d_ of > 30 mM. However, ChnR natively regulates cyclohexanol catabolism and is activated by its natural ligand, cyclohexanone, at sub-millimolar concentrations^11,12^. Yeom et al. selected mutants of the NitR biosensor to sense caprolactam at concentrations as low as 50 μM when added exogenously, and leveraged this sensor to identify novel cyclases to convert 6ACA to caprolactam^10^. Natively, NitR regulates nitrile catabolism in *Rhodococcus rhodochrous* J1, and is responsive to micromolar concentrations of isovaleronitrile^13,14^ It is reasonable to assume then that if natural lactam catabolic pathways are identified, highly sensitive biosensors could also be found.

Recently two groups have identified pathways of lactam degradation in both *P. putida* and *Pseudomonas jessenii*^15^’^16^. Work in *P. putida* demonstrated that the enzyme OplBA, putatively responsible for the hydrolysis of valerolactam, is upregulated by the lactam but not its cognate ω-amino acid^16^. These findings suggest there may be a lactam-sensitive transcription factor controlling the expression of the hydrolytic enzyme that can be used as a biosensor. In this work we demonstrate that the AraC-type regulator directly downstream of *oplBA* is indeed a lactam biosensor with unprecedented sensitivity towards both valerolactam and caprolactam. Through rational engineering we developed a suite of lactam sensing plasmids with dissociation constants ranging from 700 nM to 1.2 mM, allowing for a dramatic dynamic range of sensing. To demonstrate the utility of these sensors, we show that they are able to detect low titers of caprolactam produced biologically in an *Escherichia coli* system.

## RESULTS

### Identification and development of *oplR* as a lactam biosensor

In *P. putida*, the *oplBA* locus is flanked by the LysR-family regulator PP_3513 upstream, and the AraC-family regulator PP_3516 downstream. To infer if either of these transcription factors regulates *oplBA*, we used publicly available fitness data to assess if either regulator is cofit with *oplBA* (http://fit.genomics.lbl.gov)^17^ While no cofitness was observed between PP_3513 and either *oplA* or *oplB*, PP_3516 was highly cofit with both genes (0.91:PP_3514, 0.80:PP_3515). To examine the hypothesis that PP_3516 is the regulator of *oplBA*, we examined the genomic contexts of the oxoprolinase loci across multiple bacteria (Figure 1). While both regulators were found in closely related species, in more distantly related *Pseudomonads*, such as *Pseudomonas aeruginosa*, only the AraC-family regulator is conserved (Figure 1). Using Multiple EM for Motif Elicitation (MEME)^18^ we attempted to identify conserved putative binding sites upstream of *oplBA* as well as PP_3516. In closely related *Pseudomonads*, including *P. aeruginosa*, a conserved motif was identified upstream of both *oplBA* as well PP_3516 (Figure 1). Attempts to confirm this as the binding site of PP_3516 were hampered by the insolubility of PP_3516 when expressed heterologously (Figure S1), a common issue with AraC-family proteins^19,20^.

**Figure 1.**
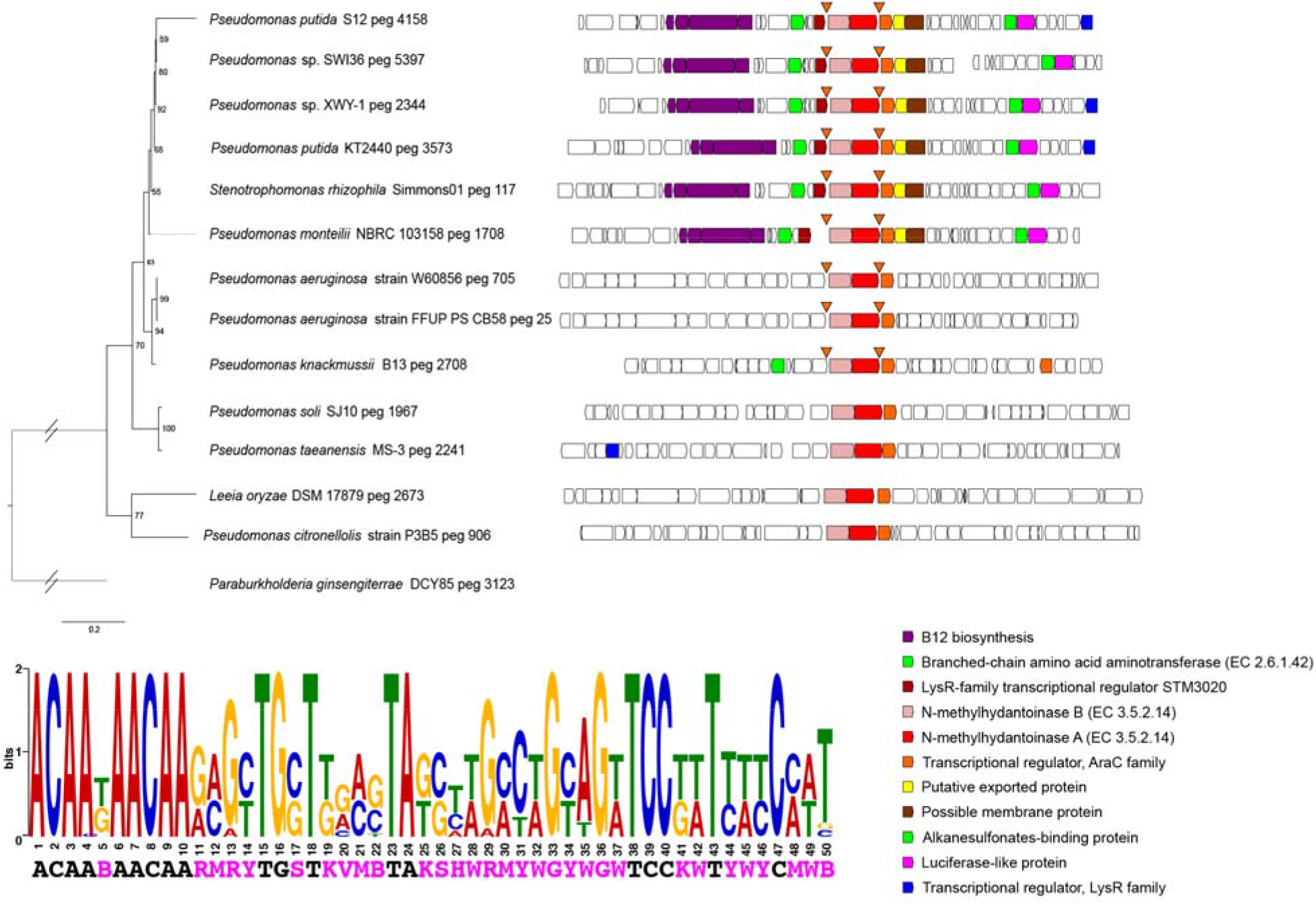
Synteny analysis of *oplBA* homologs across genomes of related species. Triangles show the location of a conserved putative binding sites of OplR. Analysis of the conserved OplR putative binding motif and its consensus sequence is shown below.

In order to screen the ability of PP_3516 to sense lactams, we employed a two-plasmid test system wherein PP_3516 was cloned into an arabinose inducible medium-copy p15a plasmid and the 200-bp upstream of *oplB* (Figure 2A) was cloned upstream of RFP on a compatible medium-copy pBBR plasmid (Figure 2B). The relationship between fluorescence output, *oplR* expression, and ligand induction was tested via a checkerboard assay where the levels of arabinose and valerolactam were varied independently of one another in cultures of *E. coli* that harbored both the “reporter” and “regulator” plasmids. A dose-dependent expression of RFP was observed; both arabinose and valerolactam were required for high-level expression of RFP (Figure 2C, Figure S2). Our initial screen also showed that the fluorescence was far above background RFP expression even at the lowest concentration of valerolactam tested (10 μM).

**Figure 2.**
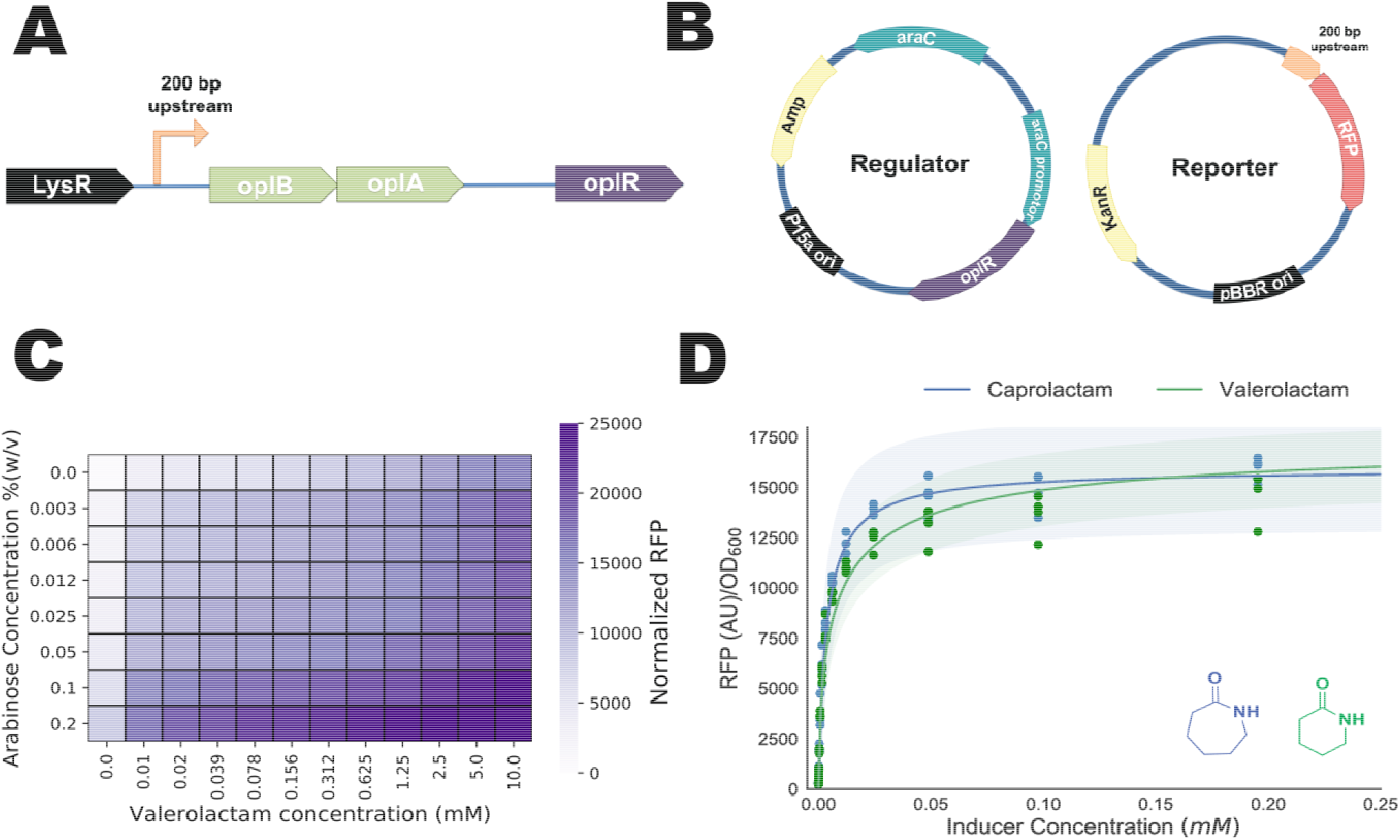
Development of an *oplR-based* lactam biosensor. A) Operonic structure of *oplBA* relative to *oplR* and putative promoter region used to construct the reporter. B) Diagram of the two-plasmid system used to test OplR lactam sensing C) Checkerboard screen of OplR biosensor two-plasmid system. Y-axis shows the concentration of arabinose (%w/v), X-axis shows the concentration of valerolactam (mM). Colorbar to right shows fluorescent intensity normalized to OD_600_, n=3. Standard error measurements are shown in Figure S2 D) Fluorescence data fit to the Hill equation to derive biosensor performance characteristics for valerolactam and caprolactam from 0 to 0.25 mM ligand. Points represent individual measurements. Shaded area represents (+/-) one standard deviation, n=4.

To quantify the sensing properties of PP_3516, the regulator was induced with a fixed concentration of arabinose at 0.0125% w/v and concentrations of either valerolactam or caprolactam were varied from 1 mM to 12 nM (Figure 2D, Figure S3). PP_3516 proved to be extremely sensitive to both caprolactam and valerolactam with both ligands having a K_d_ ~5 μM, and limits of detection ≤ 12 nM (Table 1). Based on these findings we propose PP_3516 be named *oplR* for <u>o</u>xo<u>p</u>ro<u>l</u>inase r<u>e</u>gulator, which encodes a biosensor ~5000x more sensitive towards caprolactam than the next most sensitive published biosensor (50 μM by an engineered NitR from Yeom et al.)^10^.

**Table 1.**
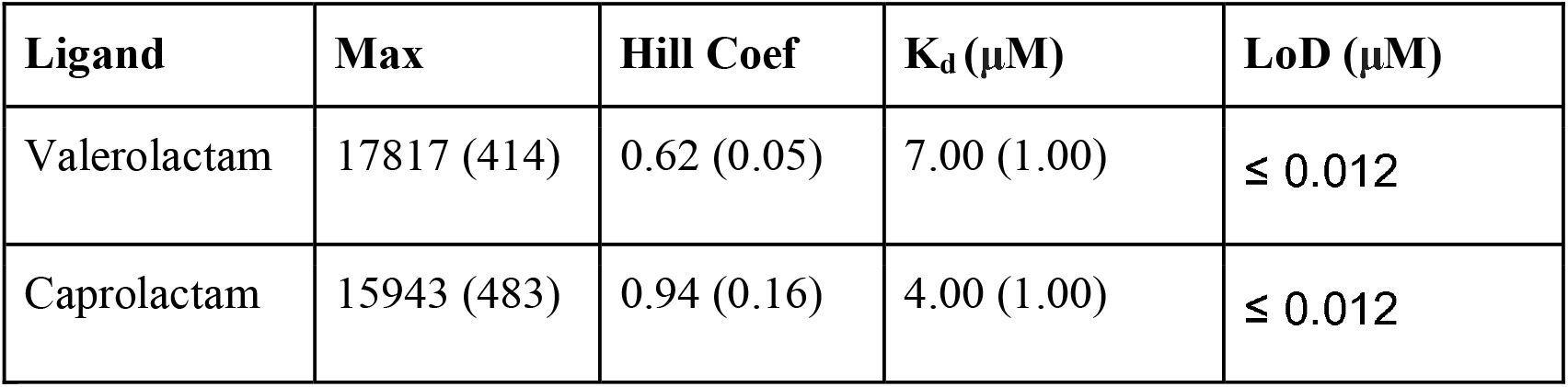
Two-plasmid biosensor parameters with caprolactam and valerolactam as ligands. Max: Predicted maximal RFP, Hill Coef: Predicted Hill coefficient, K_d_: Predicted K_d_ in μM. LoD: Limit of detection determined experimentally. Standard deviation estimates are in parentheses.

| Ligand | Max | Hill Coef | K<sub>d</sub> ($\mu\text{M}$) | LoD ($\mu\text{M}$) |
| --- | --- | --- | --- | --- |
| Valerolactam | 17817 (414) | 0.62 (0.05) | 7.00 (1.00) | $\leq 0.012$ |
| Caprolactam | 15943 (483) | 0.94 (0.16) | 4.00 (1.00) | $\leq 0.012$ |

### Development of One Plasmid Systems

As a two-plasmid system is not convenient for engineering biological systems, we then sought to consolidate both the reporter and regulator into a single vector. Initial screening of OplR in the checkerboard assay suggested that varying the level of expression of OplR could dramatically influence the resulting sensing properties of the system (Figure 2C). We therefore constructed a family of plasmids, pLACSENS, where *oplR* was constitutively expressed from five promoters of increasing strength divergent from the RFP reporter (Figure 3A).

**Figure 3.**
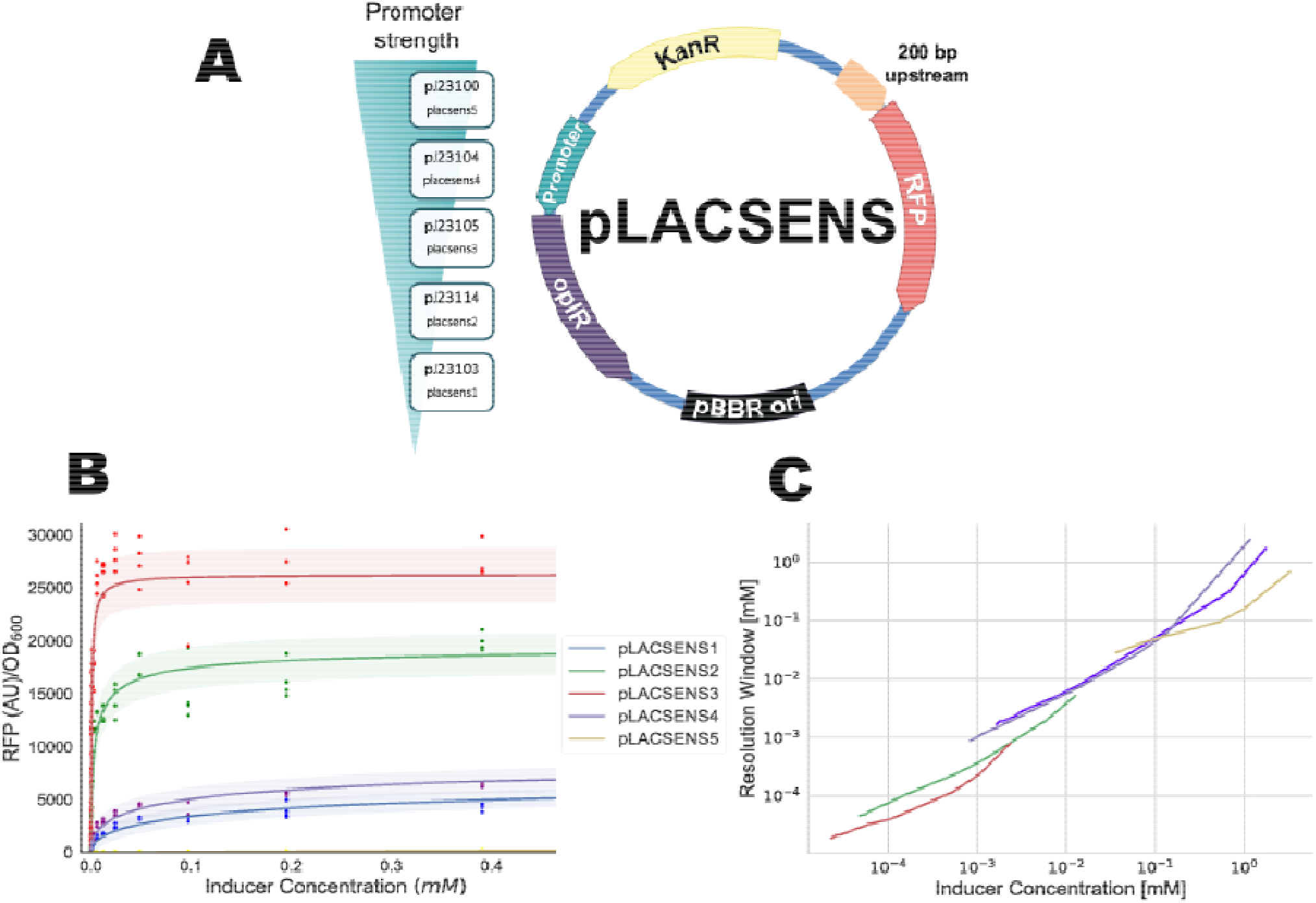
Development of pLACSENS lactam biosensors A) Diagram of the pLACSENS vector design, with the relative predicted strength of the promoter driving *oplR* on the left B) Fluorescence data fit to the Hill equation to derive biosensor performance characteristics for valerolactam against all 5 pLACSENS vectors. Points represent individual measurements. Shaded area represents (+/-) one standard deviation, n=4. C) Resolution window of each pLACSENS vector over a range of valerolactam concentrations.

Biosensor performance of pLACSENS vectors was then assessed using valerolactam as a ligand across concentrations from 12.5 mM to 12.5 nM, as the 6-membered ring of valerolactam is more stable in aqueous solution than the 7-membered ring of caprolactam (Figure 3B). As seen in the two-plasmid system, by varying the strength of *oplR* expression the characteristics of the biosensor changed dramatically (Table 2). The most sensitive vector, pLACSENS3, had an experimentally determined limit of detection (LoD) of ≤12 nM, and a K_d_ of 700 nM. The least sensitive vector, pLACSENS5, had a limit of detection of 1.5 μM, and a K_d_ of 1.5 mM, but drives the expression of *oplR* with the strongest predicted promoter. The maximal RFP expression also varied greatly with *oplR* expression, with the highest and lowest RFP expression observed in pLACSENS3 and pLACSENS5, respectively (26300 vs. 793 RFP (AU)/OD_600_). Induction over background expression was also highly variable; pLACSENS1 and pLACSENS4 were both maximally induced at ~250x over background, while pLACSENS5 was only induced ~25x over background. Time course analysis of *E. coli* harboring pLACSENS3 broadly showed that at high concentrations of valerolactam, fluorescence can be observed above background ~2.5 hours after the initiation of growth (Figure S4). Maximal RFP is achieved between 10-15 hours, depending on the concentration of ligand (Figure S4).

Given the wide range of biosensing parameters within the pLACSENS vectors we sought to characterize which ligand concentration ranges each vector is most suited to detect. To do this, we utilized a recently developed model to probabilistically relate inducer concentration and fluorescence data via Markov Chain Monte Carlo (MCMC) sampling^21^. A resolution window is defined as the concentrations of inducer that are statistically compatible with the fluorescence data fit to the Hill function at a 95% confidence interval (cI). Resolution windows for each pLACSENS plasmid were graphed from their experimentally determined LoD to ligand concentrations compatible with 75% maximal fluorescence (Figure 3C). Overall, the family of vectors showed high relative resolution from 5 nM to 1 mM valerolactam, with pLACSENS3 having the highest resolution at the lowest concentrations and pLACSENS5 having the best resolution at high concentrations (Figure 3C).

**Table 2.** One-plasmid biosensor parameters: Max: Predicted maximal RFP, Hill Coef: Predicted Hill coefficient, K_d_: Predicted K_d_ in μM, Exp. LoD in μM: Limit of detection determined experimentally. Induction: maximal induction over background based on experimental data.

| Plasmid | Max | Hill Coef | K<sub>d</sub> (μM) | LoD (μM) | Induction |
| --- | --- | --- | --- | --- | --- |
| pLACSENS1 | 8284 (358) | 0.54 (0.05) | 190 (50) | 0.024 | 253x |
| pLACSENS2 | 19614 (419) | 0.63 (0.05) | 4.00 (0.60) | ≤ 0.012 | 93x |
| pLACSENS3 | 26300 (390) | 0.90 (0.07) | 0.70 (0.10) | ≤ 0.012 | 38x |
| pLACSENS4 | 9877 (421) | 0.52 (0.05) | 90.0 (20.0) | 0.024 | 256x |
| pLACSENS5 | 793 (25) | 0.96 (0.10) | 1.23 (0.15) | 1.50 | 25x |

### OplR is selective for valerolactam and caprolactam

To assess lactam specificity of OplR, we measured fluorescence induction of pLACSENS3 in the presence of lactams (laurolactam, caprolactam, valerolactam, butyrolactam, 5-oxoproline), ω-amino acids (4-aminobutyrate, 5AVA, 6ACA), the lactone valerolactone, and piperidine (Figure 4). Robust fluorescence induction was observed with caprolactam, valerolactam, and 5AVA (Table 3), but no other tested chemicals were capable of induction at the concentrations tested in the work (data not shown).

**Figure 4.**
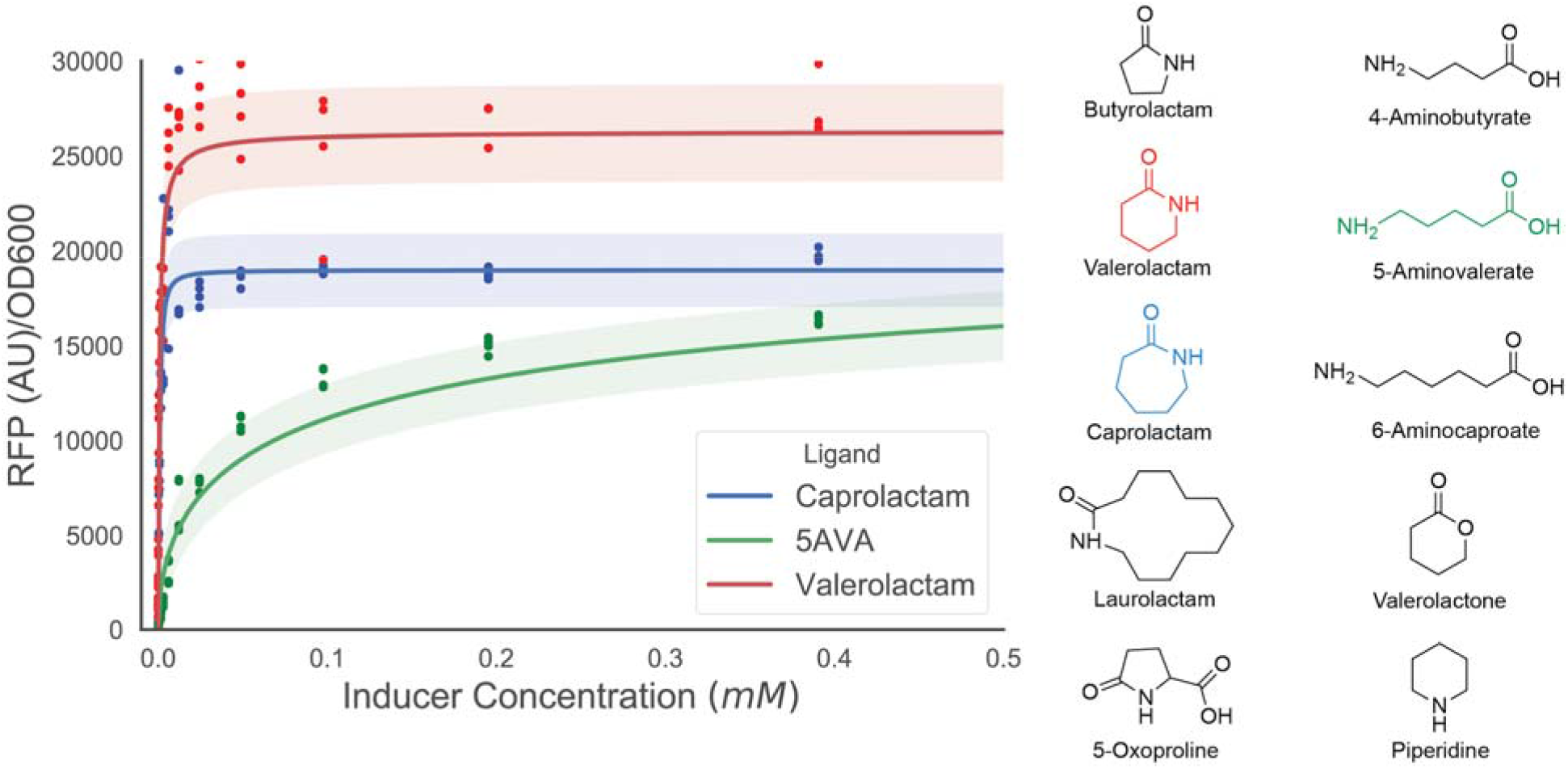
Ligand range of pLACSENS3. Fluorescence data fit to the Hill equation to derive biosensor performance characteristics for ligands that activated pLACSENS3. Points represent individual measurements. Shaded area represents (+/-) one standard deviation, n=4. To the right, chemical structures of ligands that were tested.

**Table 3.** pLACSENS3 biosensor parameters against different ligands: Max: Predicted maximal RFP, Hill Coef: Predicted Hill coefficient, K_d_: Predicted K_d_ in μM.

| Ligand | Max | Hill Coef | K<sub>d</sub> (μM) |
| --- | --- | --- | --- |
| Valerolactam | 26300 (390) | 0.90 (0.07) | 0.70 (0.10) |
| Caprolactam | 18974 (272) | 1.36 (0.14) | 0.90 (0.10) |
| 5AVA | 23766 (1093) | 0.52 (0.05) | 120 (40.4) |

Given the dissimilarity in the chemical structure of 5AVA and lactams, the ability of pLACSENS3 to detect 5AVA was surprising. Previously, it has been shown in *E. coli* that 5AVA can spontaneously be converted into valerolactam by the activity of native acyl-coA ligases^1^. This led us to believe that rather than detecting 5AVA, pLACSENS3 was detecting valerolactam derived from the added 5AVA. We fed *E. coli* cultures harboring pLACSENS3 multiple concentrations of 5AVA, measured the resulting fluorescence, and determined the final valerolactam concentrations using LC-TOF. The fluorescence signal from the spontaneously produced valerolactam was nearly identical to fluorescence signal from valerolactam supplied exogenously at the same concentration (Figure S5). These data highly suggest that 5AVA is not a ligand of OplR.

### Detection of caprolactam production *in vivo*

To demonstrate the utility of *oplR* based systems for metabolic engineering applications, we introduced our most sensitive pLACSENS plasmid (pLACSENS3) into *E. coli* harboring various acyl-coA ligases or an acyl-carrier protein sham control on IPTG-inducible orthogonal plasmids (Figure 5A). Multiple reports have utilized acyl-coA ligases to cyclize exogenously added 6ACA to produce low titers of caprolactam, with production ranging from 0.8 – 2 mg/L^1,3^. Strains harboring both plasmids were grown for 24 hours in LB medium supplemented with 10 mM, 5mM, 1mM, or 0 mM 6ACA. Cells grown in the presence of 6ACA demonstrated fluorescence greater than cells grown without 6ACA (Figure 5B). LC-MS analysis confirmed that no detectable caprolactam was produced in cells grown without 6ACA, while cells grown with 5mM and 10 mM 6ACA had produced ~6 μg/L and ~11 μg/L, respectively (Figure 5B). Although cells fed 1 mM 6ACA showed higher fluorescence than cells not fed the precursor, no caprolactam was detected above the LC-QTOF limit of detection (~1.8 μg/L). Interestingly there was no significant difference in both fluorescence or actual production between strains expressing coA-ligases versus the acyl-carrier protein control (Figure 5B). Comparison of fluorescence signal to caprolactam production in cells fed either 0 mM, 5 mM, or 10 mM showed a strong Spearman’s rank correlation (*ρ*=0.865, *p*=1.8E-18) (Figure 5C). The ability of pLACSENS3 to detect such minute production validates its utility as a means to rapidly and accurately screen novel pathways for caprolactam production.

**Figure 5.**
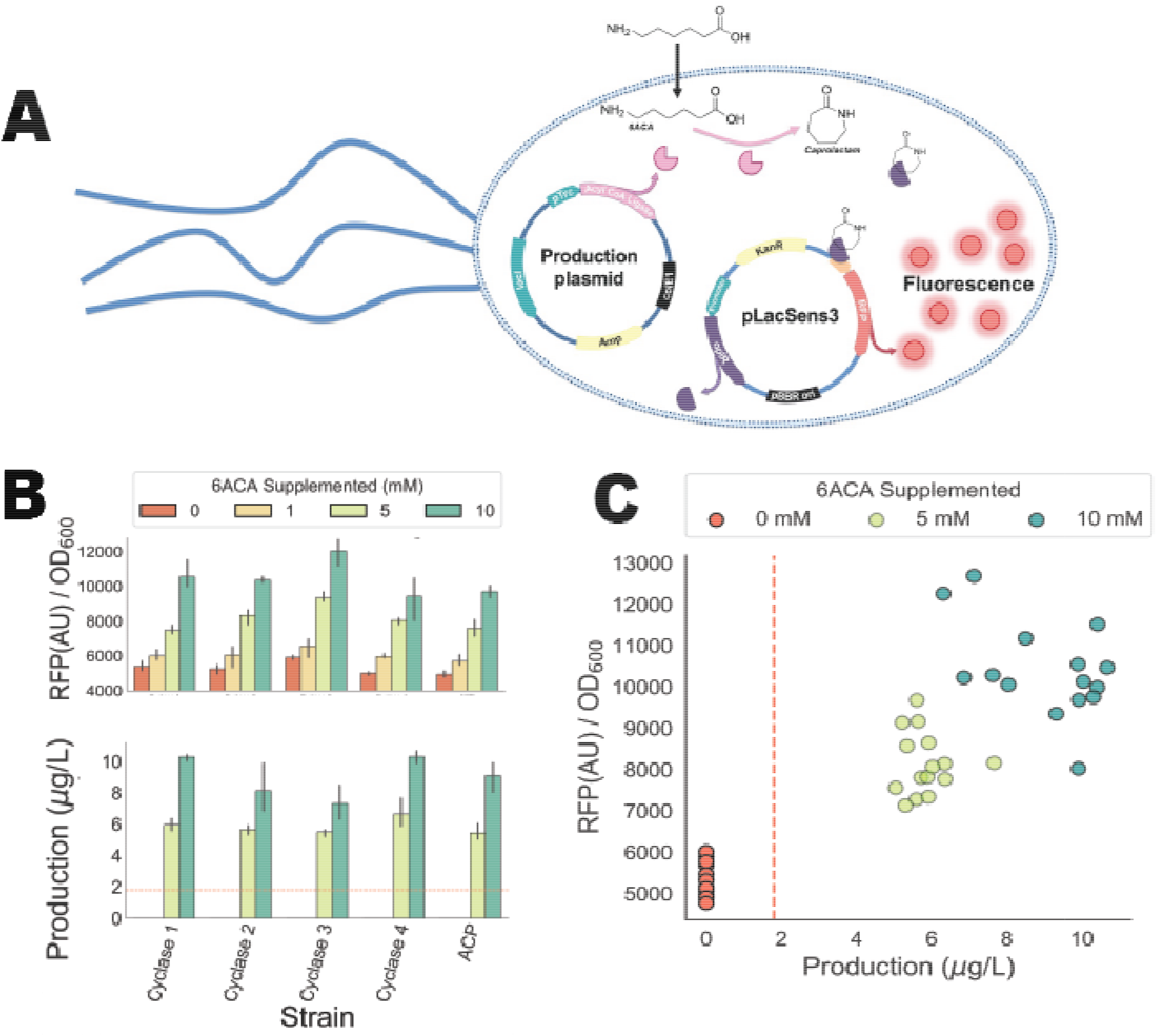
Detection of caprolactam production in *E. coli* via pLACSENS3. A) Diagram illustrating the application of the pLACSENS3 system to monitor *in vivo* caprolactam production B) Four different acyl-CoA ligases and a sham control were tested for the *in vivo* cyclization of 6ACA to caprolactam. These included the previously studied ORF26 CoA-ligase^1^ and three additional homologs. Cells were fed either 0 mM (red bars), 1 mM (yellow bars), 5 mM (green bars), or 10 mM (teal bars) 6ACA and were measured for fluorescence (top panel) or caprolactam production (bottom panel) after 24 hours, n=3. Dashed red line in the bottom panel shows the LC-MS limit of detection (~1.8 μg/L). C) Correlation of fluorescence and caprolactam production in 6ACA feeding experiment. Circle color shows amount of 6ACA supplementation. Dashed line on the x-axis shows the LC-MS limit of detection (~1.8 μg/L).

## DISCUSSION

Previously, we have leveraged shotgun proteomics to infer local regulation within the lysine catabolism of *P. putida^16,21–23^*. From these data, multiple glutarate biosensors were engineered and used to measure relative metabolite amounts in the native host^21^. Here we again leveraged previously published proteomics data that showed OplBA to be specifically upregulated in the presence of valerolactam to develop a biosensor^16^. The AraC-family regulator tentatively named OplR was shown to have limits of detection for exogenously added valerolactam or caprolactam ≤ 12 nM when expressed at particular levels. This is remarkably more sensitive than the previously published valerolactam and caprolactam sensors, which showed limits of detection between 50-100 **μ**M^10^. We attribute this high degree of sensitivity to the fact that, to our knowledge, it is the first sensor to specifically control the catabolism of lactams.

The range of detection of either caprolactam or valerolactam was highly dependent on the expression of *oplR*, with the K_d_ varying from 700 nM to 1.23 mM depending on the constitutive promoter used to drive expression. Previous work has also demonstrated that the sensing parameters of transcription factors can be readily modulated by changing the strength of transcription factor expression^24^. At both the highest and lowest levels of predicted expression, OplR was less sensitive to lactams than when *oplR* was expressed more moderately. This may be explained by the fact that some AraC-family regulators are known to act as both positive and negative regulators, thus overexpression of OplR could result in hyper-repression^20^. Unfortunately, insoluble expression of OplR prevented further examination of the biochemical means of transcriptional control.

OplR was shown to be highly specific for valerolactam and caprolactam as ligands, but not butyrolactam or laurolactam. These findings are consistent with previous observations that showed *oplBA* mutants were not defective in their ability to hydrolyze butyrolactam. The inability of *oplR* to sense the annotated substrate of OplBA, 5-oxoproline, suggests that the natural function of the amidohydrolase is not that of a 5-oxoprolinase. While the 5-membered lactam rings tested here were not able to induce OplR-mediated expression, further work should be conducted to test derivatives of valerolactam and caprolactam. Additional functional groups added to these lactams could be used to produce both pharmaceutical precursors and polymers with novel nylon properties. Identifying biological routes to their synthesis is highly attractive. Furthermore, recent work has shown that directed evolution may also be applied to broaden the ligand range^26^, which could allow OplR to accommodate different lactam ligands.

Current published routes to caprolactam production *in vivo* rely on the cyclization of 6ACA via the activity of promiscuous CoA-ligase activity^1,3^. While OplR-based biosensors may be able to aid in the selection of mutant acyl-coA ligases with enhanced activity, there remains a sizeable thermodynamic barrier for the cyclization of the 7-membered ring^1^. Novel routes to caprolactam or naturally occurring caprolactam-containing natural products that can mitigate this barrier would be ideal for high-level production. For example, a better understanding of pestalactam A-C biosynthesis may provide new and more efficient chemoenzymatic routes to 7-ring cyclization^27^ Recent work biochemically characterizing the fluvirucin β-amino acid loading pathway may also help open the door to effective PKS-based routes to non-natural lactam synthesis^28^.

In addition to the value of OplR as a biosensor for metabolic engineering purposes, it may also be useful as an inducible system. Valerolactam is inexpensive (~$2/gram), and highly water soluble (291 mg/mL). The vector pLACSENS2 demonstrated ~100x induction over background, with a K_d_ toward valerolactam of 4 uM, and the second highest maximal expression of any single vector tested. Additional engineering of the system could improve upon these qualities. Furthermore, as OplR works well in *E. coli* and is derived from the distantly related *P. putida*, it may work well in other bacterial systems. Future work could evaluate which hosts are suitable for this inducible system.

Since naturally-occuring genetically encoded biosensors for chemicals of interest have the potential to be much more sensitive than those repurposed or evolved in the laboratory, it is critical to pursue rapid and efficient means of identifying them. The recent development of other high-throughput methods to associate genotypes with phenotypes, such as RB-TnSeq and CRISPRi, has created a large reservoir of data that can be easily mined for transcription factors useful in synthetic biology^22,29–31^. Bacteria often locally regulate catabolism, thus allowing inference of genetic control by adjacent transcription factors once a catabolic pathway has been discovered. Empirical evidence of catabolism is critical for assigning transcription factor function, as orthologous transcription is often utilized differently by different species^32^. Future work to generalize approaches to develop useful synthetic biology tools from genome-wide fitness data has the potential to dramatically increase the genetically encoded chemical sensor space.

## METHODS

### Media, chemicals, and culture conditions

General *E. coli* cultures were grown in Lysogeny Broth (LB) Miller medium (BD Biosciences, USA) at 37 °C. When indicated, *E. coli* was also grown on EZ-RICH medium (Teknova, Hollister, CA) supplemented with 1% glucose. Cultures were supplemented with kanamycin (50□dmg/L, Sigma Aldrich, USA), or carbenicillin (100mg/L, Sigma Aldrich, USA), when indicated. All other compounds were purchased through Sigma Aldrich (Sigma Aldrich, USA).

### Strains and plasmids

All bacterial strains and plasmids used in this work are listed in Table 4. All strains and plasmids created in this work are available through the public instance of the JBEI registry. (https://public-registry.jbei.org/folders/XXX). All plasmids were designed using Device Editor and Vector Editor software, while all primers used for the construction of plasmids were designed using j5 software^33–35^. Plasmids were assembled via Gibson Assembly using standard protocols^36^, or Golden Gate Assembly using standard protocols^37^ Plasmids were routinely isolated using the Qiaprep Spin Miniprep kit (Qiagen, USA), and all primers were purchased from Integrated DNA Technologies (IDT, Coralville, IA).

**Table 4:** Strains and plasmids used in this study

| Strain | JBEI Part ID | Reference |
| --- | --- | --- |
| E. coli DH10B |  | 41 |
| E. coli BL21(DE3) |  | Novagen |
| P. putida KT2440 |  | ATCC 47054 |
| **Plasmids** |  |  |
| pBADT |  | 42 |
| pBADT-PP_3516p-RFP | JBEI-104519 | This work |
| pBbA8a-PP_3516 | JBEI-104517 | This work |
| pLacSens1 | JBEI-104506 | This work |
| pLacSens2 | JBEI-104505 | This work |
| pLacSens3 | JBEI-104504 | This work |
| pLacSens4 | JBEI-104503 | This work |
| pLacSens5 | JBEI-104502 | This work |
| pBbE7a-ORF26 | JBEI-104322 | This work |
| pBbE7a-G2NX2_STRVO | JBEI-104323 | This work |
| pBbE7a-A0A0D4DX08_9ACTN | JBEI-104324 | This work |
| pBbE7a-A0A0C1VDH3_9ACTN | JBEI-104325 | This work |
| pBbE7a-NmexACP-6xHis |  | 43 |

### Expression and purification of proteins

Proteins were purified as described previously^38^. The cultures were grown at 37 °C until the OD_600_ nm reached 0.8 and cooled on ice for 20 min. 1 mM IPTG was added to induce overexpression for 16 h at 18 °C. The cells were harvested by centrifugation (8000g, 10 min, 4 °C), resuspended in 40 mL of lysis buffer (50 mM HEPES, pH 8.0, 0.3 M NaCl, 10% glycerol (v/v) and 10 mM imidazole), and lysed by sonication on ice. Cellular debris was removed by centrifugation (20000g, 60 min, 4 °C).

The supernatant was applied to a fritted column containing Ni-NTA resin (Qiagen, USA) and the proteins were purified using the manufacturer’s instructions. Fractions were collected and analyzed via SDS-PAGE. Fluorescence biosensor assays

All endpoint assays were conducted in 96-deep well plates (Corning Costar, 3960), with each well containing 500 **μ**L of medium with appropriate ligands, antibiotics, and/or inducers inoculated at 1% v/v from overnight cultures. Plates were sealed with AeraSeal film (Excel Scientific, AC1201-02 and incubated at 37 C in a 250 rpm shaker rack. After 24 hours, 100 **μ**L from each well was aliquoted into a black, clear-bottom 96-well plate for measurements of optical density and fluorescence using an Infinite F200 (Tecan Life Sciences, San Jose, CA) plate reader. Optical density was measured at 600 nm (OD_600_), while fluorescence was measured using an excitation wavelength of 535 nm, an emission wavelength of 620 nm, and a manually set gain of 60. To perform time course assays, overnight cultures were inoculated into 10 mL of LB medium from single colonies, and grown at 37□°C. These cultures were then diluted 1:100 into 100 uL of EZ-Rich medium with 1% (w/v) glucose in 96-well plates (Falcon, 353072). Plates were sealed with a gas-permeable microplate adhesive film (VWR, USA), and then optical density and fluorescence were monitored for 48 hours in a Tecan Infinite F200 (Tecan Life Sciences, San Jose, CA) plate reader. Optical density was measured at 600 nm (OD_600_), while fluorescence was measured using an excitation wavelength of 535 nm, an emission wavelength of 620 nm, and a manually set gain of 60.

For the checkerboard assay of the two-plasmid system, LB medium supplemented with both kanamycin and carbenicillin was inoculated with E. coli containing the regulator and reporter plasmids grown overnight in the same medium. Arabinose concentration was decreased from 0.2 to 0% w/v along the y-axis, while valerolactam concentration was increased from 0-10 mM along the x-axis.

To find the Hill fit to the two-plasmid system, EZ-RICH medium containing kanamycin, carbenicillin, and 0.0125 w/v% arabinose was inoculated with an overnight culture of the two-plasmid system in LB medium supplemented with both kanamycin and carbenicillin. Both valerolactam and caprolactam were tested at concentrations ranging from 0 to 50 mM.

Characterization of the 5 variations of the one-plasmid pLACSENS plasmids was conducted using EZ-rich medium containing kanamycin and inoculated with overnight cultures of the appropriate E. coli strain. Valerolactam concentrations was varied from 0 to 50 mM. This same assay was repeated on pLACSENS3 with various ligands—using concentrations between 0 to 50 mM of caprolactam and 5AVA, and 0 to 10 mM of butyrolactam, laurolactam, 5-oxoproline, gamma-aminobutyric acid, 6ACA, valerolactone, and piperidine.

### Production assays and analytical methods

Caprolactam production assays were carried out in 10 mL of LB medium supplemented with 10 mM 6ACA, as well as kanamycin, carbenicillin, and 1 mM IPTG. Cultures were inoculated 1:100 with overnight culture harboring both pLACSENS3 and expression vectors for acyl-coA ligases and grown at 30 °C shaking at 200 rpm for 24 hours. After 24 hours optical density was measured at 600 nm (OD_600_), while fluorescence was measured using an excitation wavelength of 535 nm, an emission wavelength of 620 nm, and a manually set gain of 60. To sample for caprolactam production 200 **μ**L of culture was quenched with an equal volume of icecold methanol and then stored at −20 °C until analysis. The different acyl-CoA ligases tested for the *in vivo* cyclization of 6ACA to caprolactam included the previously studied ORF26 CoA-ligase (cyclase 1) and three additional homologs. Cyclase 2 refers to A0A0D4DX08_9ACTN, cyclase 3 refers to G2NX2_STRVO and cyclase 4 refers to A0A0C1VDH3_9ACTN. An acyl-carrier protein was used as a sham control.

Valerolactam and caprolactam were measured via LC-QTOF-MS as described previously^16^. Liquid chromatographic separation was conducted at 20°C with a Kinetex HILIC column (50-mm length, 4.6-mm internal diameter, 2.6-μm particle size; Phenomenex, Torrance, CA) using a 1260 Series HPLC system (Agilent Technologies, Santa Clara, CA, USA). The injection volume for each measurement was 5 μL. The mobile phase was composed of 10 mM ammonium formate and 0.07% formic acid in water (solvent A) and 10 mM ammonium formate and 0.07% formic acid in 90% acetonitrile and 10% water (solvent B) (HPLC grade, Honeywell Burdick & Jackson, CA, USA). High purity ammonium formate and formic acid (98-100% chemical purity) were purchased from Sigma-Aldrich, St. Louis, MO, USA. Lactams were separated with the following gradient: decreased from 90%B to 70%B in 2 min, held at 70%B for 0.75 min, decreased from 70%B to 40%B in 0.25 min, held at 40%B for 1.25 min, increased from 40%B to 90%B for 0.25 min, held at 90%B for 1 min. The flow rate was varied as follows: 0.6 mL/min for 3.25 min, increased from 0.6 mL/min to 1 mL/min in 0.25 min, and held at 1 mL/min for 2 min. The total run time was 5.5 min.

The HPLC system was coupled to an Agilent Technologies 6520 quadrupole time-of-flight mass spectrometer (QTOF MS) with a 1:6 post-column split. Nitrogen gas was used as both the nebulizing and drying gas to facilitate the production of gas-phase ions. The drying and nebulizing gases were set to 12 L/min and 30 lb/in^2^, respectively, and a drying gas temperature of 350°C was used throughout. Fragmentor, skimmer and OCT 1 RF voltages were set to 100 V, 50 V and 300 V, respectively. Electrospray ionization (ESI) was conducted in the positive-ion mode for the detection of [M + H]^+^ ions with a capillary voltage of 4000 V. The collision energy voltage was set to 0 V.

MS experiments were carried out in the full-scan mode (75–1100 *m z)* at 0.86 spectra/s. The QTOF-MS system was tuned with the Agilent ESI-L Low concentration tuning mix in the range of 50-1700 *m/z.* Lactams were quantified by comparison with 8-point calibration curves of authentic chemical standards from 0.78125 μM to 100 μM. R^2^ coefficients of ≥0.99[EB1] were achieved for the calibration curves. Data acquisition was performed by Agilent MassHunter Workstation (version B.05.00), qualitative assessment by Agilent MassHunter Qualitative Analysis (version B.05.00 or B.06.00), and data curation by Agilent Profinder (version B.08.00) Bioinformatic Analysis

For the phylogenetic reconstructions, the best amino acid substitution model was selected using ModelFinder as implemented on IQ-tree phylogenetic trees were constructed using IQ-tree, nodes were supported with 10,000 bootstrap replicates. The final tree figures were edited using FigTree v1.4.3 (http://tree.bio.ed.ac.uk/software/figtree/). Orthologous syntenic regions of OplBA were identified with CORASON-BGC^40^ and manually colored and annotated. DNA-binding sites were predicted with MEME^18^.

### Supporting Information

The Supporting Information is available free of charge on the ACS Publications website. Supporting information contains figures that show 1) Expression of OplR 2) Standard errors of checkerboard assays 3) Fluorescent data across all concentrations tested in two-plasmid system 4) Time course fluorescence data of *E. coli* harboring pLACSENS3 5) Comparison of fluorescence signal from valerolactam produced by feeding 5AVA to valerolactam added directly. The SI also contains detailed methods on analyzing biosensor performance.

## Acknowledgements

The views and opinions of authors expressed herein do not necessarily state or reflect those of the United States Government or any agency thereof. Neither the United States Government nor any agency thereof, nor any of their employees, makes any warranty, expressed or implied, or assumes any legal liability or responsibility for the accuracy, completeness, or usefulness of any information, apparatus, product, or process disclosed, or represents that its use would not infringe privately owned rights. The United States Government retains and the publisher, by accepting the article for publication, acknowledges that the United States Government retains a nonexclusive, paid-up, irrevocable, worldwide license to publish or reproduce the published form of this manuscript, or allow others to do so, for United States Government purposes. The Department of Energy will provide public access to these results of federally sponsored research in accordance with the DOE Public Access Plan (http://energy.gov/downloads/doe-public-access-plan).

## Contributions

Conceptualization, M.G.T.; Methodology, M.G.T.,A.N.P.,J.F.B, P.C.M.,E.E.K.B.,N.S,Z.C.; Investigation, M.G.T., L.E.V., J.F.B, P.C.M, M.R.I.,M.E.G.,A.N.P.,; Writing – Original Draft, M.G.T.; Writing – Review and Editing, All authors.; Resources and supervision, H.G.M,A.M.,J.D.K.

