## Supplementary Info for "Identification, characterization, and application of a highly sensitive lactam biosensor from *Pseudomonas putida*"

**Table of Contents**

**Supporting Figures**

1. **Figure S1 - Expression of OplR**
2. **Figure S2 - Standard Error of Checkerboard Assays**
3. **Figure S3 - Two-plasmid fluorescent data across all concentrations tested**
4. **Figure S4 - Time course fluorescence data of *E. coli* harboring pLACSENS3**
5. **Figure S5 - Comparison of fluorescence signal from valerolactam produced by feeding 5AVA to valerolactam added directly**

**Supporting Methods**

1. **Analysis of Biosensor Performance**

**Supporting References**

**Supporting Figures**

**Figure S1: Insolubility of OplR expressed heterologously in *E. coli*: OplR-6xHis expressed in the insoluble fraction (~40 kD) of *E. coli* in the presence and absence of the detergent Tween 20 (0.5% w/v).**

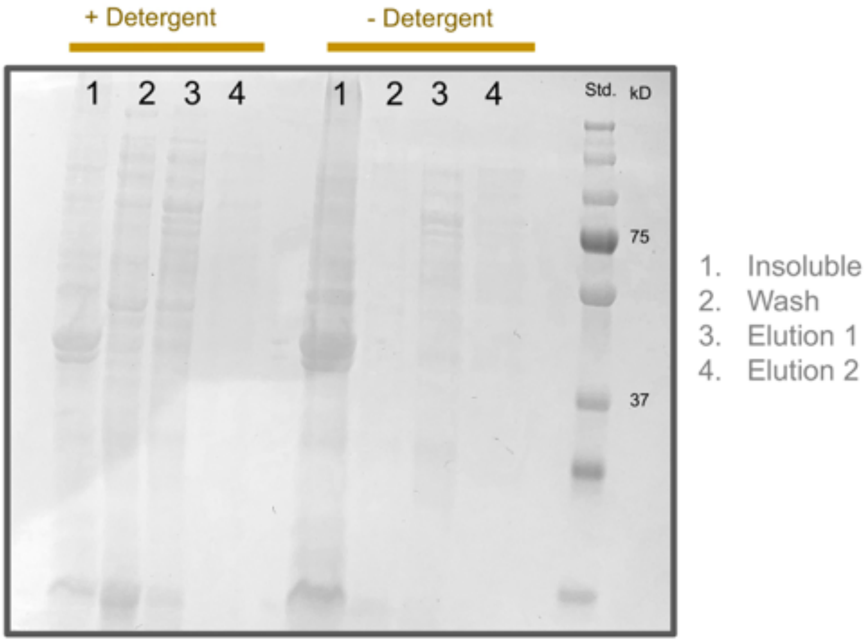

**Figure S2: Standard error of checkerboard screen of OplR biosensor two-plasmid system. Y-axis shows the concentration of arabinose (%w/v), X-axis shows the concentration of valerolactam (mM). Colorbar to right shows fluorescent intensity normalized to $OD_{600}$ (n=3)**

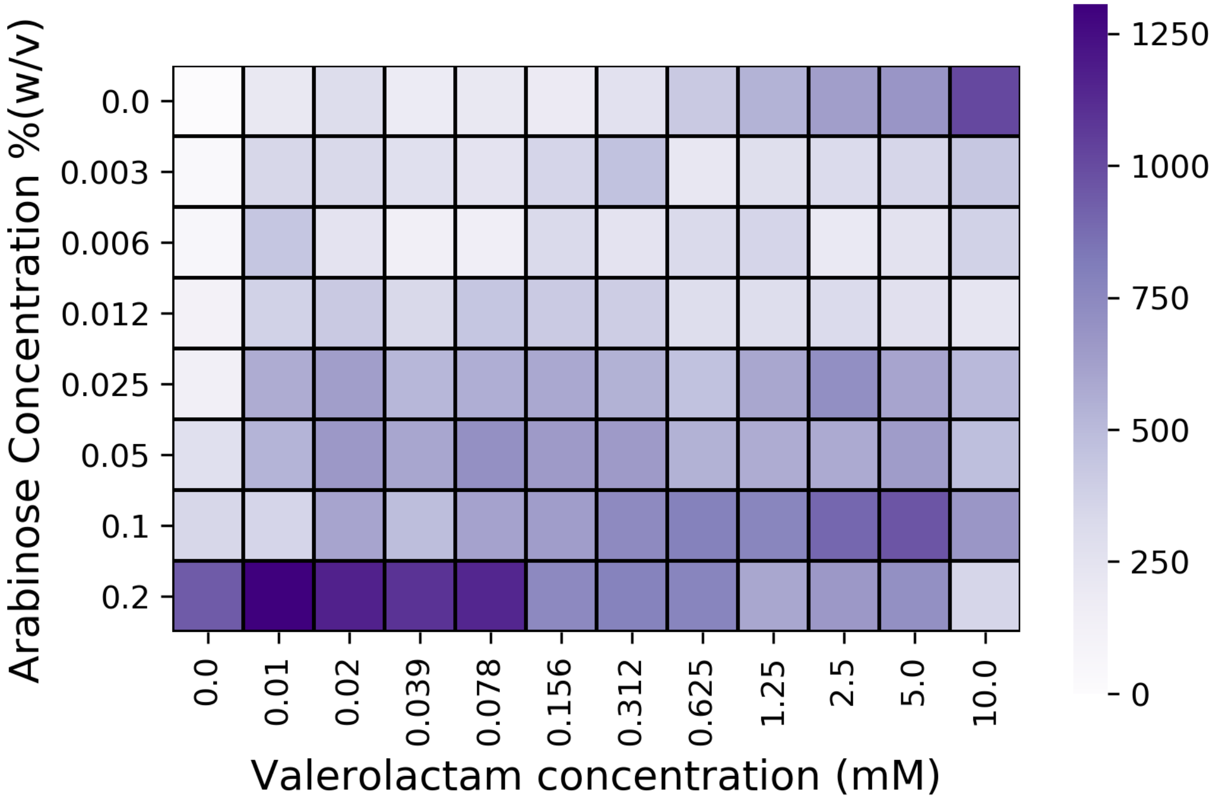

**Figure S3: Fluorescence data fit to the Hill equation to derive biosensor performance characteristics for valerolactam and caprolactam from 0 to 12.5 mM ligand. Points represent individual measurements. Shaded area represents (+/-) one standard deviation, n=4.**

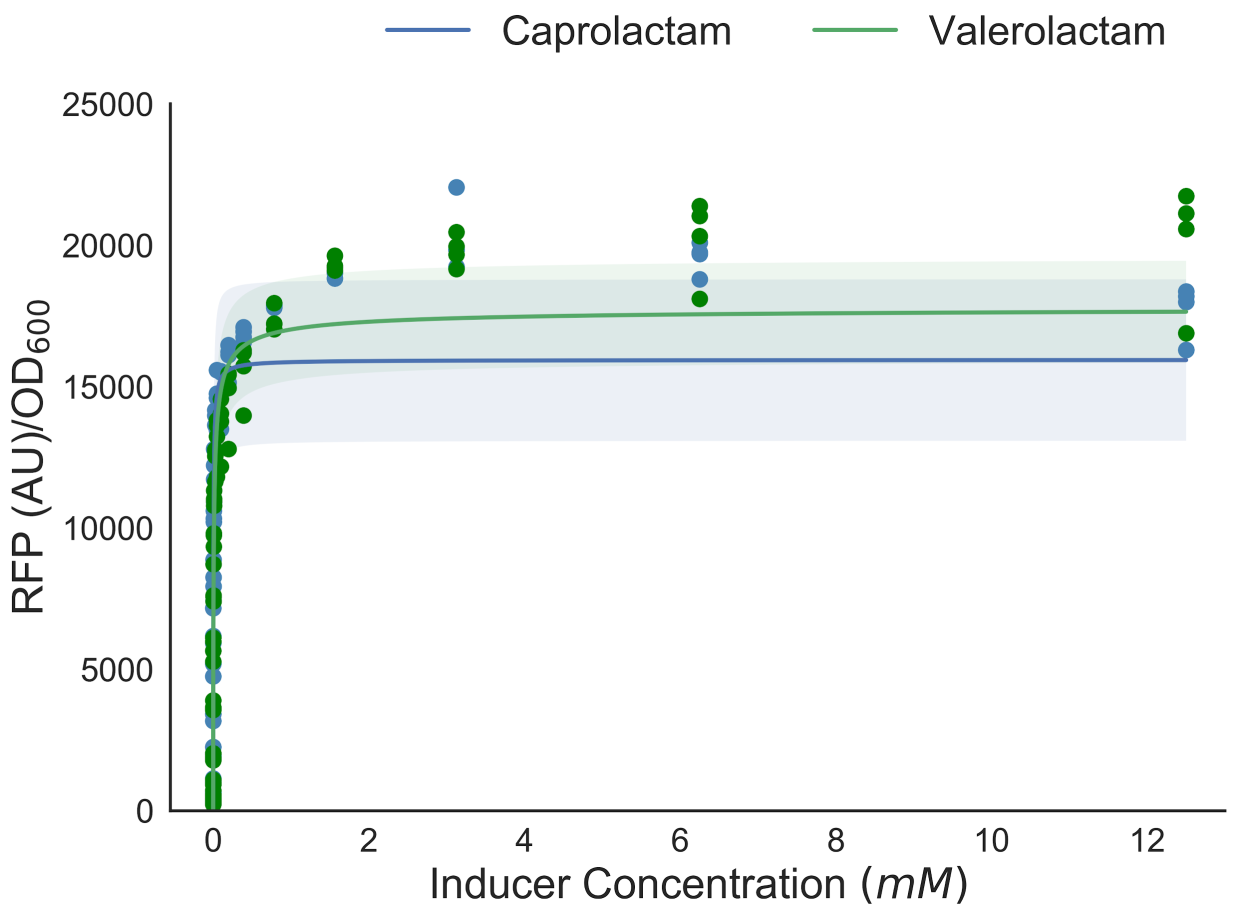

**Figure S4: Time course fluorescence data of *E. coli* harboring pLACSENS3 from 0 to 100 µM valerolactam. The above graph shows data from 0 to 24 hours, while the lower graph shows a subset of data from 2 to 5 hours highlighted by the red box. Shaded area represents (+/-) one standard deviation, n=8.**

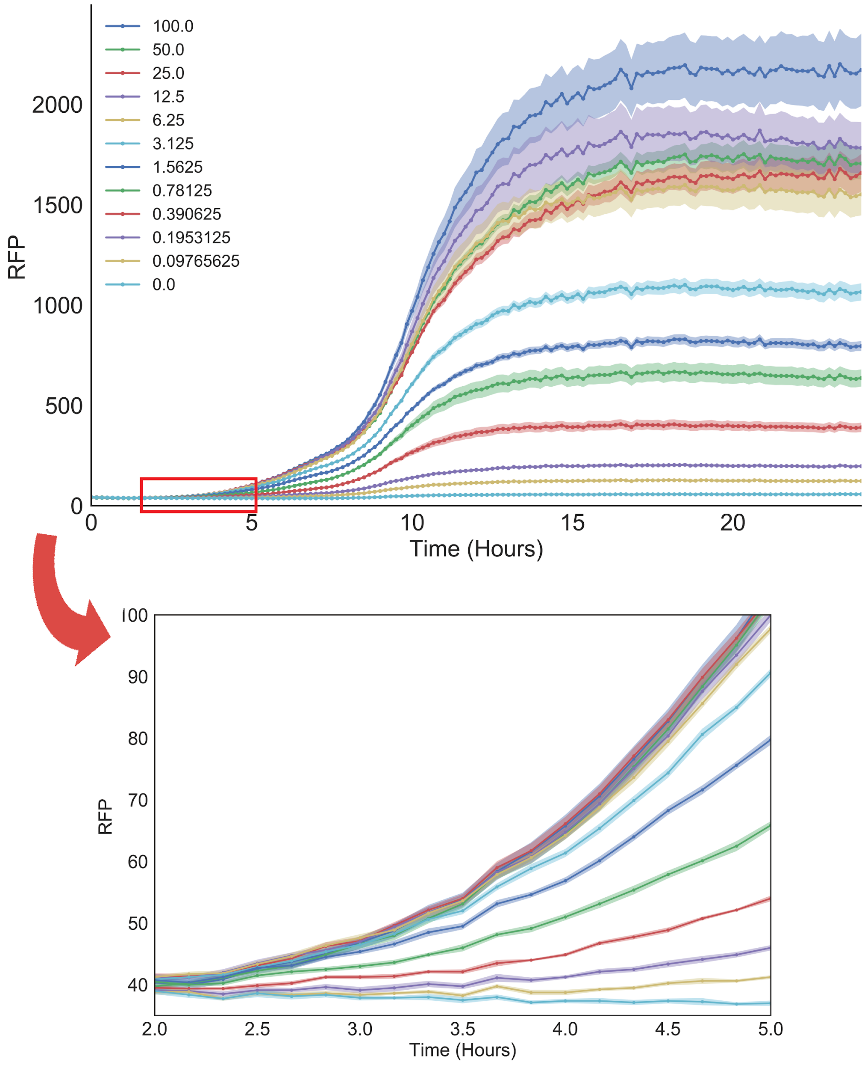

**Figure S5: Comparison of fluorescence signal from valerolactam produced by feeding 5AVA to valerolactam added directly. Black dots show fluorescence signal from *E. coli* harboring pLACSENS3 after 24-hours. Colored dots show fluorescence signal and valerolactam present as measured by LC-MS when fed 1000 µM (gold), 62 µM (red), or 31µM (blue) 5AVA for 24 hours.**

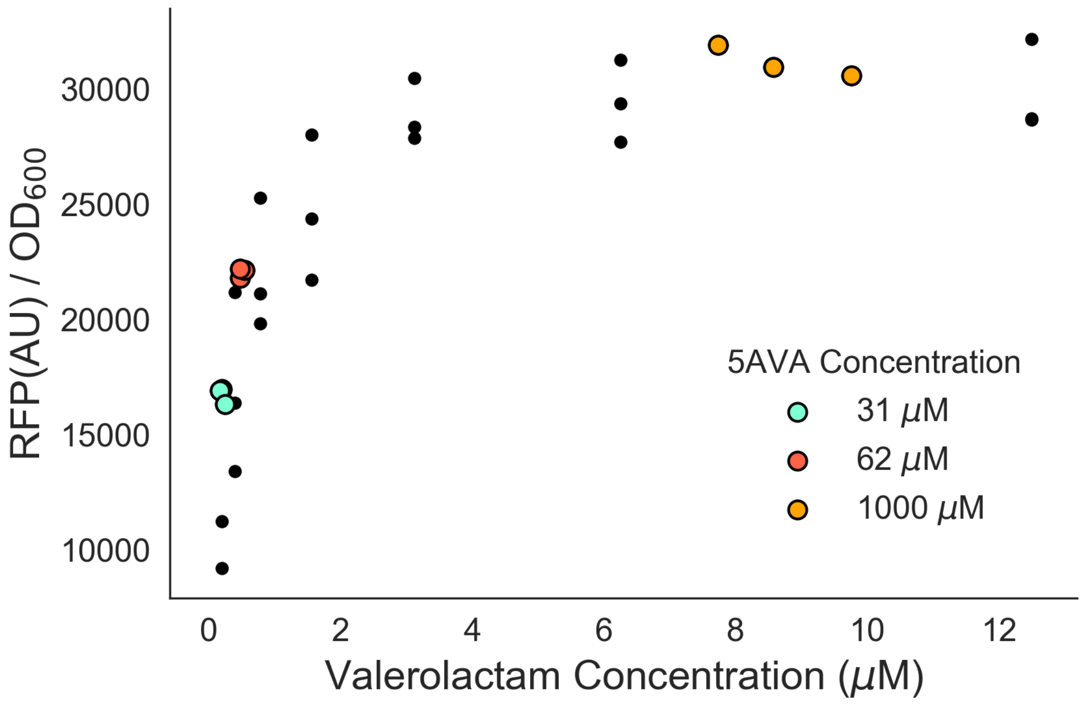

### Supporting Methods

Analysis of Biosensor Performance

Fluorescent measurements were fit to the Hill equation and biosensor parameters we estimated as described previously (Thompson et al. 2019).

We determined biosensor resolution by solving the above maximum likelihood estimation problem iteratively over the range of observed fluorescences during the biosensor characterization process. This can determine the relationship between an inducer concentration estimate and the estimated standard deviation. The standard deviation of the estimate of inducer concentration can be interpreted as the resolution window. Here, two standard deviations is considered the resolution window of the sensor.

Induction above background was calculated by dividing the maximal experimental normalized RFP expression by background fluorescence in uninduced cultures. The experimental limit of detection was defined as the minimal concentration of inducer that produced normalized fluorescence that was statistically above uninduced cultures harboring the same plasmid via Student's t-test ($p<0.05$) .
